# Significant organic carbon acquisition by *Prochlorococcus* in the oceans

**DOI:** 10.1101/2022.01.14.476346

**Authors:** Zhen Wu, Dikla Aharonovich, Dalit Roth-Rosenberg, Osnat Weissberg, Tal Luzzatto-Knaan, Angela Vogts, Luca Zoccarato, Falk Eigemann, Hans-Peter Grossart, Maren Voss, Michael J. Follows, Daniel Sher

**Affiliations:** Department of Earth, Atmospheric and Planetary Sciences, Massachusetts Institute of Technology, Cambridge, MA 02139, USA; Department of Marine Biology, Leon H. Charney School of Marine Sciences, University of Haifa, 31905, Israel; Leibniz-Institute for Baltic Sea Research, Seestrasse 15, D-18119 Warnemuende, Germany; Department of Experimental Limnology, Leibniz-Institute of Freshwater Ecology and Inland Fisheries, Alte Fischerhuette 2, D-16775 Stechlin, Germany; Potsdam University, Institute of Biochemistry and Biology, Maulbeeralle 2, D-14469 Potsdam, Germany

## Abstract

Marine phytoplankton are responsible for about half of the photosynthesis on Earth. Many are mixotrophs, combining photosynthesis with heterotrophic assimilation of organic carbon but the relative contribution of these two carbon sources is not well quantified. Here, single-cell measurements reveal that *Prochlorococcus* at the base of the photic zone in the Eastern Mediterranean Sea are obtaining only ~20% of carbon required for growth by photosynthesis. Consistently, laboratory-calibrated evaluations of *Prochlorococcus* photosynthesis indicate that carbon fixation is systematically too low to support published *in situ* growth rates in the deep photic layer of the Pacific Ocean. Furthermore, agent-based model simulations show that mixotrophic cells maintain realistic growth rates and populations 10s of meters deeper than obligate photo-autotrophs, deepening the nutricline and Deep Chlorophyll Maximum by ~20 m. Time-series of *Prochlorococcus* ecotype-abundance from the subtropical North Atlantic and North Pacific suggest that up to 30% of the *Prochlorococcus* cells live where light intensity is not enough to sustain obligate photo-autotrophic populations during warm, stratified periods. Together, these data and models suggest that mixotrophy underpins the ecological success of a large fraction of the global *Prochlorococcus* population and its collective genetic diversity.

---

Photosynthesis by phytoplankton provides most of the energy and fixed carbon that support marine food webs and carbon reservoirs^1^. However, few phytoplankton are strictly photo-autotrophic^2^. Many phytoplankton also utilize dissolved organic matter, taking up particulate detrital organic matter or preying upon other living cells and even harvesting organelles^2^. Mixotrophic lifestyles, in which cells both fix carbon and use exogenously available organic carbon, may enhance fitness when the relative availability of inorganic resources differs from physiological demands^3^. This may occur, for example, where light intensity is low but inorganic nutrients are abundant. Despite the potential importance of mixotrophy to phytoplankton life history, the contribution of heterotrophic carbon assimilation to phytoplankton growth is not well quantified^4^. Simulations suggest that mixotrophy may be a globally significant carbon source for phytoplankton^5^ but this prediction is currently difficult to quantitatively test with empirical data. One reason is that dissolved organic carbon (DOC) in the oceans constitutes an extremely complex mixture of compounds^6,7^, most of which are uncharacterized. This means that uptake measurements using specific organic carbon sources (e.g. glucose, amino acids)^8,9^ do not represent the entirely available DOC pool and may underestimate the actual DOC uptake rates, and hence mixotrophy of major phytoplankton species^10^.

*Prochlorococcus* are the most abundant phototrophic cells on Earth, actively growing at depths ranging from the ocean surface down to the base of the photic zone (~160 m)^11^. Across these depths, photosynthetically available radiation (PAR) varies over 3-4 orders of magnitude, a challenge which the diverse *Prochlorococcus* lineage faces using a variety of adaptations in their photosynthetic apparatus^11,12^. These adaptations have led to the diversification of *Prochlorococcus* into high-light and low-light adapted clades^11,12^. In addition, *Prochlorococcus* are mixotrophs, able to uptake dissolved organic compounds such as glucose^8^, pyruvate^13^, amino acids^9^, nucleotides^10^ and perhaps DMSP^14,15^. Yet, to what extent DOC uptake can supplement or replace photosynthetically fixed carbon for respiration and/or growth in this globally-abundant lineage is still unknown^10^. Available evidence suggests that while mixotrophy helps *Prochlorococcus* survive limited periods of darkness, axenic cells die after ~1 week if not exposed to light^13,16^ indicating that light harvesting, and possibly photosynthesis, are likely obligate.

Here, we take a multi-faceted approach to evaluate the contribution of heterotrophic carbon assimilation to *Prochlorococcus* in the oceans. We first use isotopic measurements to quantify photosynthesis and N uptakes in wild *Prochlorococcus* populations at the base of the photic zone in the Mediterranean Sea. Then we compare observed growth rates from the Pacific Ocean with purely photo-autotrophic growth rates simulated by a laboratory-calibrated photo-physiological model. We also use an individual-based model to illustrate how mixotrophy provides a fitness advantage and deepens the nutricline. Finally, we use time-series observations of vertical profiles of *Prochlorococcus* ecotypes in subtropical gyres to show that several clades rely extensively on mixotrophic carbon assimilation. Overall, these results suggest that up to a quarter of depth integrated carbon assimilation by *Prochlorococcus* originates from DOC, with implications for global C cycles, and that mixotrophy is essential to support a significant fraction of *Prochlorococcus* diversity.

## Results and discussion

### Carbon and nitrogen uptake in wild samples from the base of the photic zone

To evaluate the relative contributions of photosynthesis and heterotrophic carbon uptake in a natural *Prochlorococcus* population from the base of the photic zone, where light may be limiting, we assess the *Prochlorococcus* population structure and per-cell activity during late summer in the ultra-oligotrophic Eastern Mediterranean Sea^17^. At the time of sampling, the water column was highly stratified, nutrients were depleted down to around 140 m, and a prominent Deep Chlorophyll Maximum (DCM) was observed at depth of ~115 m (Figure 1A). *Prochlorococcus* were the numerically dominant phytoplankton below the surface (Fig 1B), and could be divided into two populations based on the per-cell fluorescence – a low fluorescence population from the surface to 115 m and a high fluorescence population from 115-150 m, with an overlap at 115 m (Figure 1C, D). The shift in the per-cell chlorophyll fluorescence in *Prochlorococcus* with depth is commonly observed^18–20^, and is usually attributed to a change in the genetic composition of the population, from High-Light adapted cells (HL, low fluorescence) to Low-Light adapted (LL, high fluorescence) ones^19^. However, phenotypic heterogeneity (acclimation) can also contribute to this phenomenon^21^, and indeed amplicon sequencing of the Internal Transcribed Spacer between the 16S and 23S genes (ITS)^21,22^ revealed a gradual transition from HL to LL clades around the DCM, suggesting both genotypic and phenotypic shifts with depth (Figure 1C). The flow cytometry and genetic data are both consistent with previous studies^21,23^, and suggest that the water column had been relatively stable for at least 3-4 days prior to sampling^20^. Notably, the light intensity at the DCM (~3-5 *μmol photons m*^−2^ *s*^−1^ during the afternoon, Figure 1A) is potentially enough under laboratory conditions to support the growth of some LL strains but not sufficient for active growth of most HL strains^24^. Since HL cells comprise >50% of the *Prochlorococcus* population at 115 m and about 25% at 125 m, this suggests that a significant fraction of the *Prochlorococcus* cells in these samples are living under conditions where photosynthesis cannot support growth (Figure 1C).

**Figure 1:**
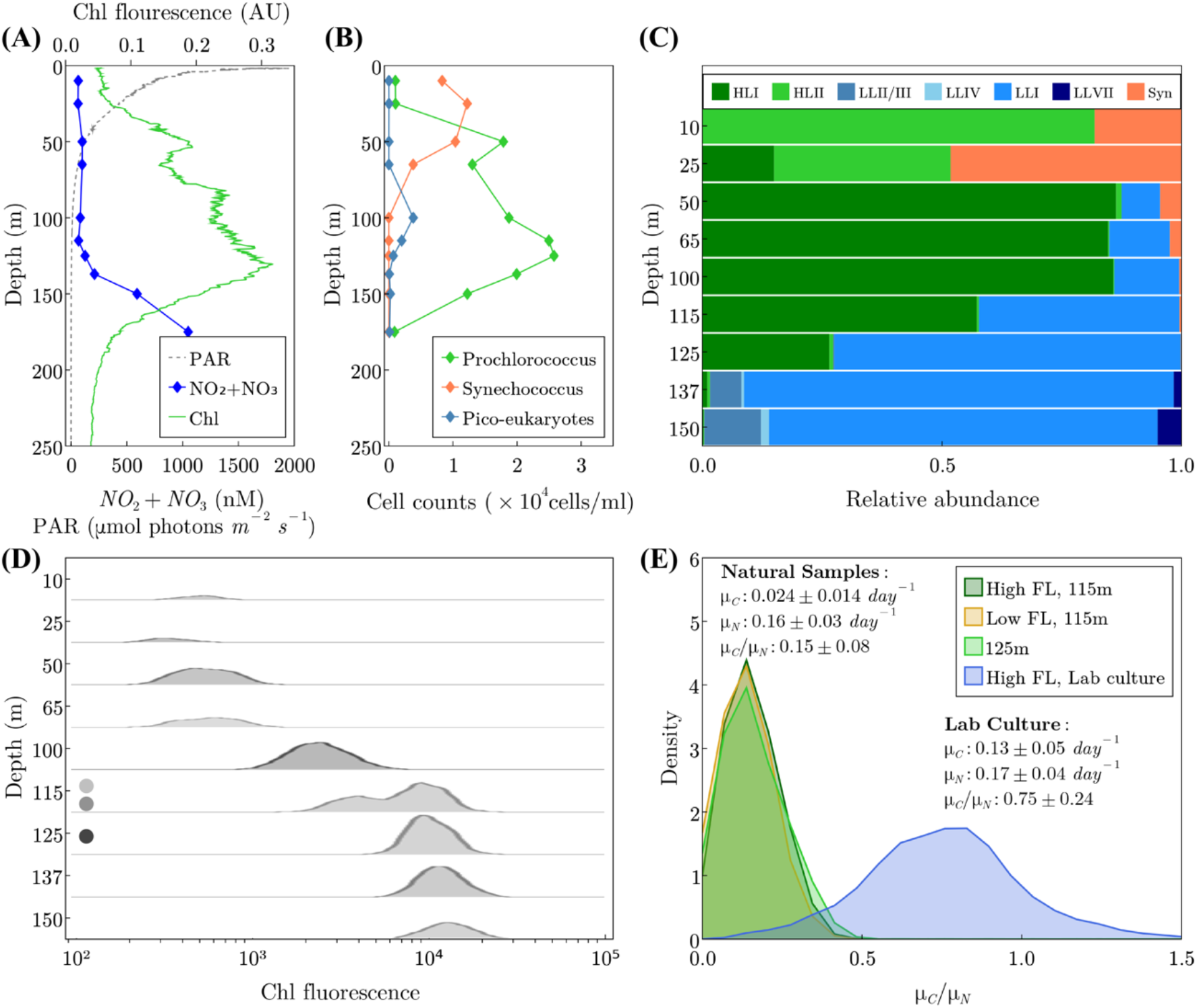
Nutrient uptake of naturally occurring *Prochlorococcus* populations at the Eastern Mediterranean Sea. (A) Depth profiles of Photosynthetically Available Radiation (PAR), NO2+NO3 and Chlorophyll. (B) Phytoplankton cell counts using flow cytometry. (C) Relative abundance of different *Prochlorococcus* clades across the water column, determined by ITS sequencing. Note the change in Chl fluorescence without a concomitant change in population structure between 60 to 100 m, as well as the presence of LL clades above 115 m and HL clades at 125 m. (D) Density plots of *Prochlorococcus* per-cell chlorophyll fluorescence. Note the double population at 115 m. The circles represent the populations sorted and analyzed by nanoSIMS. (E) Density plots of the ratios of C-specific C uptake rate (*μ_C_*) to N-specific N uptake rate (*μ_N_*) from NanoSIMS analysis of each sorted sub-population from 115 m, the single population from 125 m, and lab cultures. The numbers of cells measured in each population are 45 (LL 115m), 49 (HL 115m), 55 (125m), and 489 (lab culture).

We next measured the uptake of ^13^C-labelled bicarbonate (representing C-fixation through photosynthesis) and of ^15^N-labeled ammonium (representing nitrogen uptake) in single *Prochlorococcus* cells from the DCM, using Nanoscale Secondary Ion Mass Spectrometry (NanoSIMS). Essentially all of the *Prochlorococcus* cells at 115 and 125 m depth were active (photosynthesized and took up NH4). The observation that essentially all of the *Prochlorococcus* cells in natural samples are active is consistent with a similar study in the North Pacific^25^, and suggests that dead or chlorotic cells observed in laboratory cultures^13,26^ may be relatively rare in nature, at least during midday at the DCM. Nevertheless, the per-cell photosynthesis rates at these depths were not sufficient to support the growth rates indicated by the nitrogen-specific nitrogen uptake rates, even though the uptake experiments were performed when light intensity was maximal (Figure 1E). Previous studies from multiple oceanic regions based on cell cycle analysis and on ^14^C incorporation into divinyl-chlorophyll indicate that *Prochlorococcus* cells at depths of 100-150 m replicate every 4-7 days (a growth rate of 0.14-0.25 *day*^−1^)^27–30^. However, the observed C-specific C uptake rate (*μ_C_*) was only ~0.024 *day*^−1^, too low to support these expected growth rates, while the observed N-specific N uptake rate (*μ_N_*) was ~0.16 *day*^−1^ indicating a doubling time of ~6 days. Furthermore, *μ_C_/μ_N_* was only ~0.15 in the field, much lower than normal cells which are expected to be ~1 (*μ_C_* ≈ *μ_N_*). Indeed, *μ_C_/μ_N_* in lab cultured *Prochlorococcus* was ~0.75 (Figure 1E). Taken together, these quantitative observations suggest that >80% of the C required for the expected growth rate of these *Prochlorococcus* cells at the DCM must come from non-photosynthetic sources.

### Evaluation of potential growth rate profiles

Our Mediterranean samples suggest that a large fraction of carbon assimilated by *Prochlorococcus* in the deeper reaches of the photic zone is of organic origin. By comparing measured profiles of growth rates and modeling photosynthetic carbon fixation from sites in the Pacific, we ask if this is consistent in other regions and infers the water column integrated contribution of heterotrophy. Vaulot et al.^31^ and Liu et al.^32^ reported vertical profiles of *Prochlorococcus* division rates based on cell-cycle analysis in the Equatorial Pacific (EqPac, 0°N, 140°W) and North Pacific Subtropical Gyre (HOT, 22°45’N, 158°W; Station ALOHA), respectively. These data were obtained in the context of an extensive biogeochemical survey (JGOFS EqPac)^33^ and time-series station (HOT)^34^ and are associated with rich contextual data sets including observations of cell counts, photon fluxes and nutrient concentrations (Figure 2A, C). Calibrated by observed, noon-time PAR profiles, we simulated the daily cycle of photosynthesis and the vertical profiles of *Prochlorococcus’* carbon-specific, net photosynthesis rate (*day*^−1^). We simulated both HL and LL ecotypes, using laboratory calibrations of the photosynthesis-irradiance relationship from Moore and Chisholm^24^. Similarly, using allometric scaling for fixed-nitrogen, phosphate and dissolved iron uptake rates^35,36^, along with observed environmental concentrations, we evaluated the nutrient-specific uptake rates (*day*^−1^). Full details are presented in Materials and Methods. The estimated, purely autotrophic growth rates were determined by the most limiting resource at each depth (Figure 2B, D). Light and carbon fixation strongly limited the simulated autotrophic growth in the deeper region of the photic layer, while fixed nitrogen (HOT), iron (EqPac) and carbon fixation, due to photo-inhibition, were important near the surface (Figure 2). While the observed growth rates at the surface were mostly within the ranges predicted from the photophysiological parameters of HL and LL strains (blue shade in Figure 2B and D), the model failed to resolve the observed growth rates below ~75-100 m at both stations. Rather, the model unequivocally suggests that photosynthesis alone cannot account for the observed division rates at depth. We interpret the differences between the modeled autotrophic and observed actual growth rates at depth (red shading) to infer the minimal rate of organic carbon assimilation of *Prochlorococcus.* The two stations represent very different physical and biogeochemical regimes, yet show similar qualitative structure. Mixotrophy appears to become significant at different depths at the two stations (95 m at HOT and 60 m at EqPac) but at similar level of PAR (~ 15 *μmol photons m*^−2^ *s*^−1^, ~5% of surface PAR). Using observed cell densities^31,32^and assumed cellular carbon quotas^37^ we estimated the vertically integrated autotrophic net primary production for *Prochlorococcus* to be ~0.35 *gC m*^−2^*day*^−1^ at HOT and ~0.20 *gC m*^−2^*day*^−1^ at EqPac, with vertically integrated heterotrophic contributions (based on the red shading in Figures 2B and D) of ~0.075 *gC m*^−2^*day*^−1^ at HOT and ~0.069 *g C m*^−2^*day*^−1^ at EqPac. In other words, assimilation of organic carbon is inferred to support ~18% of total *Prochlorococcus* biomass production at HOT and ~25% at EqPac. Furthermore, organic carbon uptake contributes ~80% at HOT and 54% at EqPac of the total production below the depth where the contribution of mixotrophy is greater than photosynthesis, broadly consistent with the isotopic inference from the deep photic zone in the Mediterranean. We note that this model does not take into account exudation of organic carbon by *Prochlorococcus* which is not well constrained experimentally and would likely reduce the inferred growth rates at the surface^38–41^. Indeed, mixotrophy (uptake of glucose and amino acids) has been observed in surface *Prochlorococcus*^9,10^, suggesting that our estimate provides a lower bound of the contribution of mixotrophy to integrated *Prochlorococcus* production.

**Figure 2.**
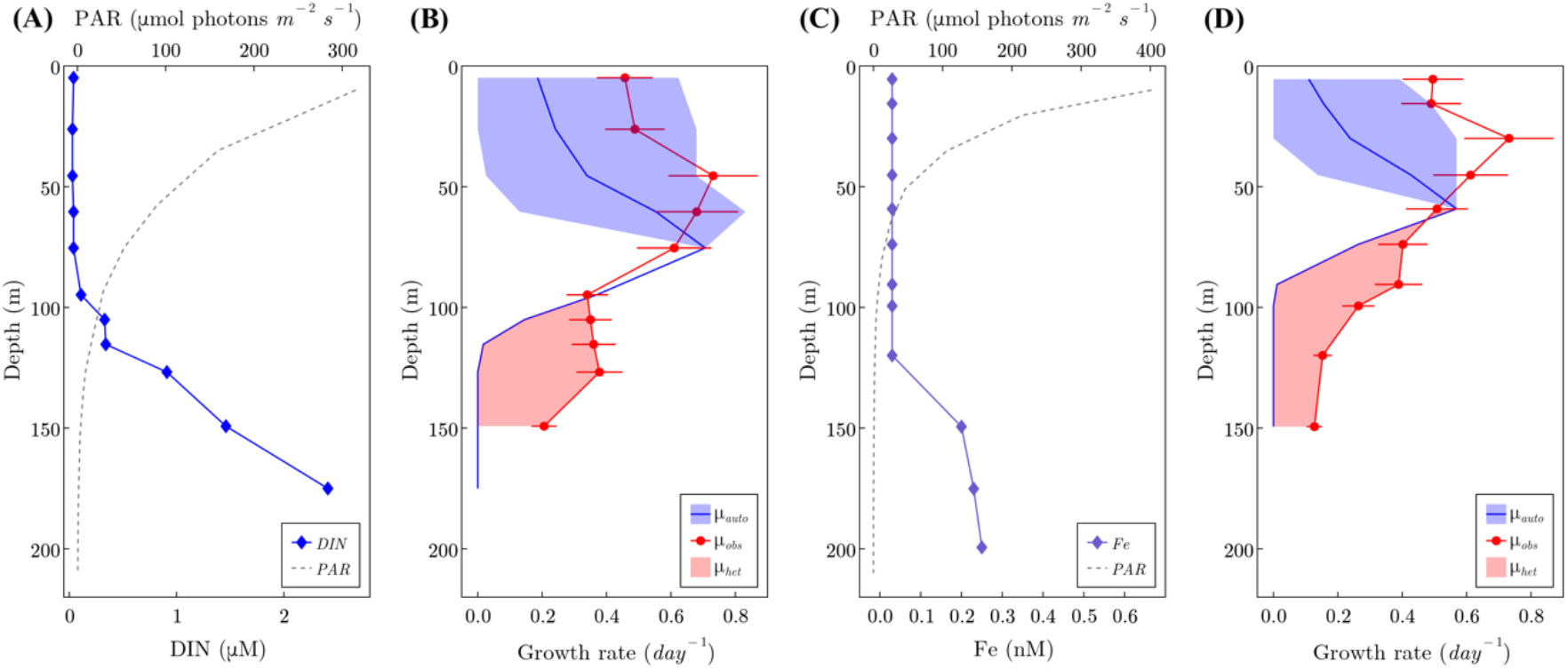
Simulated growth rates at HOT and EqPac. (A), (C) Observations of PAR and dissolved inorganic nitrogen (DIN) at the Hawaii Ocean Time-series (HOT, panel A) and of PAR and dissolved iron (Fe) at the equatorial Pacific (EqPac, panel C). (B), (D) Simulated autotrophic growth rate (blue line) and observed growth rates (red line with dots, data from^42^) at HOT (B) and EqPac (D). The blue shade represents the difference between simulated HL and LL ecotypes, and the red shade represents the inferred heterotrophic growth rate. A 19% error of observed growth rate is included both at HOT and EqPac according to Vaulot et al.^31^.

### Simulations in a dynamic water column

To investigate the implications of mixotrophy on biogeochemical dynamics, we employed an individual-based modeling approach (see Method for details), simulating trajectories of individual *Prochlorococcus* cells (or super-agents representing many cells) through light and nutrient environments in a two-dimensional, highly resolved turbulent fluid flow (see supporting movie). Inorganic nutrients and a DOC-like tracer are represented by density-based equations. Briefly, individuals fix carbon by photosynthesis and take up inorganic nitrogen and phosphorus. Two idealized types of individuals are simulated separately, one with a strict photo-autotrophic lifestyle and the other which is mixotrophic and able to assimilate carbon from the DOC-like substance. The mixotrophic individual cannot live strictly heterotrophically, as suggested by Coe et al.^13^, which we parameterize as requiring 1% of the incorporated C to come from photosynthesis. In Figure 3A we illustrate horizontally-averaged profiles of cell density from the purely autotrophic and mixotrophic simulations, illustrating how mixotrophy supports a population of *Prochlorococcus* below ~75 m. The simulated daily division rate of ~0.2 *day*^−1^ at depth (Figure 3B) is consistent with the published cell-cycle profiles from the subtropical and the Equatorial Pacific^31,32^and is a bit higher than the aforementioned inferred division rate in the Mediterranean based on NH4 uptake. Mixotrophs and autotrophs share the same division rate (~0.3 *day*^−1^) in the mixed layer (surface 50 m) where the inorganic nutrient is the limiting factor in the simulations. The autotrophs then reach a maximum daily division rate of ~0.5 *day*^−1^ at 60 m depth where the transition of N to C limitation happens, and then decrease rapidly to zero at 90 m depth due to light limitation. In contrast, the mixotrophs have a deeper maximum growth rate of ~0.5 *day* ^−1^ at 80 m depth where the transition of N to C limitation occurs and gradually decrease to ~0.2 *day*^−1^ at 125 m depth (Figure 3B). The deeper maximum division depth of the mixotrophs and their ability to maintain a population at depths where photosynthesis is not sufficient are supported by the DOC utilization, which is presented as a black line in Figure 3B. In the mixotrophic simulation, the contribution of DOC uptake to the vertically integrated total production is ~12%; ~43% when light becomes the limiting factor, below the red stripe in Figure 3B. The contribution of DOC and the maximal depth at which *Prochlorococcus* can grow are broadly consistent with the division rate profile model and are sensitive to parameter values which control the nutritional value of the DOC-like substance (and which cannot be *a priori* constrained by empirical data at this point; see Materials and Methods). Notably, the horizontal stripes in Figure 3B indicate the depth at which limitation shifted from nutrients to C in the two ensembles of simulations. This horizon is deeper when the cells are mixotrophic and leads to a significantly deeper nutricline in the simulation with mixotrophic cells (Figure 3C).

**Figure 3.**
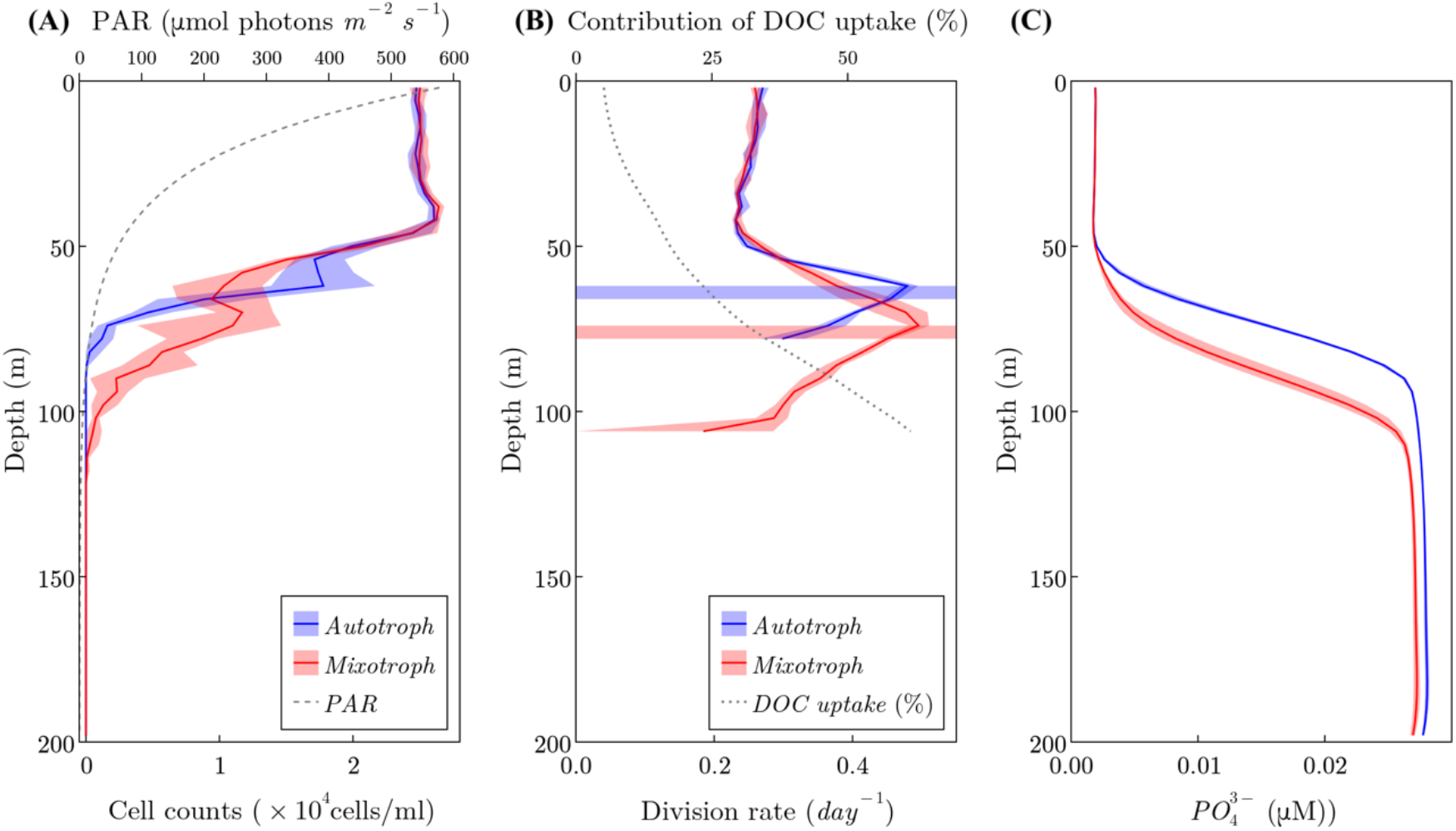
Vertical profiles of simulated autotrophs and mixotrophs in the individual-based model. The red and blue shades in all panels indicate the differences between an ensemble of 10 model runs. (A) Vertical profiles of cell density of simulated autotrophs (blue) and mixotrophs (red). The vertical profile of PAR is represented as the gray dashed line. (B) Vertical profiles of cell division rate of autotrophs (blue) and mixotrophs (red). The blue and red stripes indicate the transition point from nutrient limitation to carbon limitation of phytoplankton growth. The black dotted line represents the contributions of DOC uptake to total carbon acquisition in mixotrophs. (C) Vertical profiles of phosphate in simulations of autotroph (blue) and mixotroph (red).

### Interpretation of vertical distributions of Prochlorococcus ecotype at time-series stations

To what extent does mixotrophy supports natural, genetically-diverse, populations of *Prochlorococcus?* To answer this question, we calculated the fraction of the *Prochlorococcus* cells and of individual ecotypes living below the depth where they can be supported by photosynthesis alone over a 5-year time series in the north Atlantic and Pacific gyres (Hawaii and Bermuda time series study sites, respectively^23^). We consider only the time of the year when the water column is stratified (white regions in Figure 4), defined here as a mixed layer depth that is shallower than the photic depth (light intensity is >10 *μmol photons m*^−2^ *s*^−1^ for high-light strains or > 2.8 *μmol photons m*^−2^ *s*^−1^ for low-light strains, experimentally-determined minimal light requirement for active growth of high-light and low-light adapted strains during a 14:10 day-night cycle^24^). This is because at other times cells below the photic depth but still within the upper mixed layer could be transferred closer to the surface and therefore receive increased light. An average of ~8-10% of the *Prochlorococcus* cells during these stratified periods are likely to be light-starved (Figure 4 A, B), including the vast majority of LL adapted ecotypes (Figure 4C, D).

**Figure 4:**
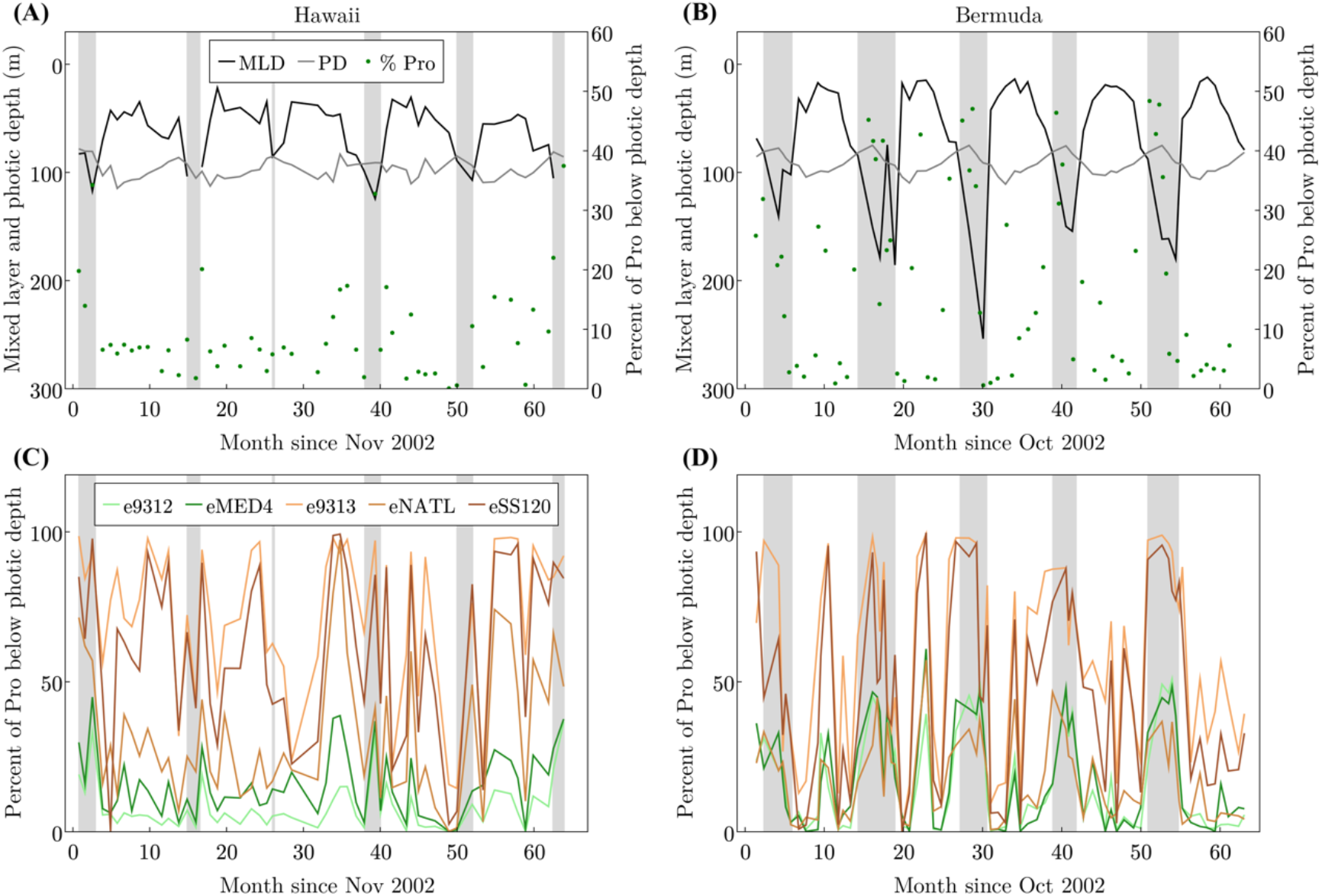
Estimating the number of *Prochlorococcus* cells and of specific ecotypes found below their photic depth at Hawaii and Bermuda. (A), (B) The percent of total *Prochlorococcus* cells (Pro) found below their photic zone at Hawaii (A) and Bermuda (B), defined as the integrated illumination level supporting the growth of representative strains in laboratory cultures^24^ (grey line shows this depth for HL strains). The black line shows the mixed layer depth (MLD), the grey line shows the photic depth (PD), the green dots represent the percentages of *Prochlorococcus* below photic depth, the grey areas are non-stratified conditions where cells may be mixed from depth to the surface. (C), (D) The percentage of each *Prochlorococcus* ecotype below its photic depth. The data are taken from Malmstrom et al.^23^.

## Conclusions

We have presented several lines of evidence illustrating the importance of mixotrophic carbon assimilation by *Prochlorococcus.* The uptake of isotopically labelled nitrogen uptake in samples from the Mediterranean Sea indicate doubling times at the DCM of about a week, consistent with cell-cycle based observations from the Equatorial and Subtropical Pacific^27,29–32^. The associated uptake of labelled carbon suggests that this growth rate is only viable if more than three-quarters of assimilated carbon is sourced from organic matter. Using a laboratory-calibrated model of carbon-specific photosynthesis rates and local environmental data, we compared carbon-limited growth rates with observed cell-cycle observations at the Pacific locations. We estimated that 18-25% of depth integrated, net carbon assimilation by *Prochlorococcus* is heterotrophic at those sites, with as much as 80% heterotrophic carbon supply at the DCM. We note that while this shifts perception of the photo-autotrophic nature of primary producers, products such as remote-sensing based estimates of global-scale primary production are typically calibrated with data from isotopically labeled inorganic carbon studies and hence, other sources of error notwithstanding, are appropriately estimating photosynthesis and not growth rates. We explored the wider consequences of the phenomenon in simulations with an individual-based model that resolves a DOC-like substance. These simulations suggest that such extensive mixotrophy in the deeper photic layer will significantly deepen the nutricline. This is significant for carbon cycle simulations, most of which do not currently resolve mixotrophy and may predict, or inappropriately tune, a too-shallow nutricline. Finally, investigation of the ecotypic, vertical biogeography in the subtropical North Pacific and North Atlantic^23^ indicates that low-light adapted *Prochlorococcus* spend 50-100% of their time, depending on season, below the deepest horizon for photo-autotrophically viable maintenance of the population. We propose that reliance on mixotrophy, rather than on photosynthesis, underpins the ecological success of a large fraction of the global *Prochlorococcus* population and its collective genetic diversity.

## Materials and Methods

### Isotope labelling and phylogenetic analysis of a natural marine bacterioplankton population at sea

Mediterranean seawater was collected during August 2017 (station N1200, 32.45°N, 34.37°E) from 11 depths by Niskin bottles and divided into triplicate 250 ml polycarbonate bottles. Two bottles from each depth were labeled with 1mM Sodium bicarbonate-^13^C and 1mM Ammonium-^15^N chloride (Sigma-Aldrich, USA) and all 3 bottles (2 labelled and 1 control) were incubated at the original depth and station at sea for 3.5 hours around mid-day. The short incubation time was chosen to minimize isotope dilution and potential recycling and transfer of ^13^C and ^15^N between community members^25^. After incubation, bottles were brought back on board and the incubations were stopped by fixing with 2X EM grade glutaraldehyde (2.5% final concentration) and stored at 4 °C until sorting analysis. Cell sorting, NanoSIMS analyses and the calculation of uptake rates were performed as described in Roth-Rosenberg et al.^26^.

### DNA collection and extraction from seawater

Samples for DNA were collected on 0.22 μm Sterivex filters (Millipore). Excess water was removed using a syringe, 1 ml Lysis buffer (40 mM EDTA, 50 mM Tris pH 8.3, 0.75 M sucrose) was added and both ends of the filter were closed with parafilm. Samples were kept at −80°C until extraction. DNA was extracted by using a semi-automated protocol including manual chemical cell lysis before the automated steps. The manual protocol began with thawing the samples, then the storage buffer was removed using a syringe and 170 μl lysis buffer added to the filters. 30 μl of Lysozyme (20 mg/ml) were added to the filters and incubated at 37°C for 30 min. After incubation, 20 μl proteinase K and 200 μl buffer AL were added to the tube for 1 hour at 56°C (with agitation). The supernatant was transferred to a new tube and DNA was extracted using the QIAcube automated system and QIAamp DNA Mini Protocol: DNA Purification from Blood or Body Fluids (Spin Protocol, starting from step 6, at the BioRap unit, Faculty of Medicine, Technion). All DNA samples were eluted in 100 μl DNA free distilled-water.

### ITS PCR amplification

PCR amplification of the ITS was carried out with specific primers for *Prochlorococcus* CS1_16S_1247F (5’-ACACTGACGACATGGTTCTACACGTACTACAATGCTACGG) and Cs2_ITS_Ar (5’-TACGGTAGCAGAGACTTGGTCTGGACCTCACCCTTATCAGGG)^21,22^. The first PCR was performed in triplicate in a total volume of 25 μl containing 0.5 ng of template, 12.5 μl of MyTaq Red Mix (Bioline) and 0.5 μl of 10 μM of each primer. The amplification conditions comprised steps at 95°C for 5 min, 28/25 (16S/ITS) cycles at 95°C for 30 sec, 50°C for 30 sec and 72°C for 1 min followed by one step of 5 min at 72°C. All PCR products were validated on a 1% agarose gel and triplicates were pooled. Subsequently, a second PCR amplification was performed to prepare libraries. These were pooled and after a quality control sequenced (2×250 paired-end reads) using an Illumina MiSeq sequencer. Library preparation and pooling were performed at the DNA Services (DNAS) facility, Research Resources Center (RRC), University of Illinois at Chicago (UIC). MiSeq sequencing was performed at the W.M. Keck Center for Comparative and Functional Genomics at the University of Illinois at Urbana-Champaign (UIUC).

### ITS Sequence processing

Paired-end reads were analyzed using the Dada2 pipeline^43^. The quality of the sequences per sample was examined using the Dada2 ‘plotQualityProfile’ command. Quality filtering was performed using the Dada2 ‘filterAndTrim’ command with parameters for quality filtering truncLen=c(290,260), maxN=0, maxEE=c(2,2), truncQ=2, rm.phix=TRUE, trimLeft=c(20,20). Following error estimation and dereplication, the Dada2 algorithm was used to correct sequences. Merging of the forward and reverse reads was done with minimum overlap of 4 bp. Detection and removal of suspected chimeras was done with command ‘removeBimeraDenovo’. In total, 388,417 sequences in 484 amplicon sequence variants (ASVs) were counted. The ASVs were aligned in MEGA6^44^ and the first ~295 nucleotides, corresponding to the 16S gene, were trimmed. The ITS sequences were then classified using BLAST against a custom database of ITS sequences from cultured *Prochlorococcus* and *Synechococcus* strains as well as from uncultured HL and LL clades.

### Individual-based Model

PlanktonIndividuals.jl (v0.1.9) was used to run the individual-based simulations. A full documentation is available at https://juliaocean.github.io/PlanktonIndividuals.jl/dev/. Briefly, the cells fix inorganic carbon through photosynthesis and nitrogen, phosphorus and DOC from the water column and grow until division or grazing. Cell division is modeled as a probabilistic function of cell size. Grazing is represented by a quadratic probabilistic function of cell population. Cells consume nutrient resources which are represented as Eulerian, density-based tracers. We set up two separate simulations, each of them either has a population of an obligate photo-autotroph or a mixotroph which also consumes DOC. The initial conditions and parameters are the same for the two simulations except the ability of mixotrophy. The simulations were run with a time step of 1 minute for 360 simulated days to achieve a steady state. We run the two simulations for multiple times in order to get the range of the stochastic processes. The code of this configuration is available at https://github.com/zhenwu0728/Prochlorococcus_Mixotrophy.

### Evaluation of autotrophic growth rates

We evaluated the carbon-specific, daily-averaged carbon fixation rate, 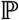 as a function of light intensity (*I, μE*) as follows:

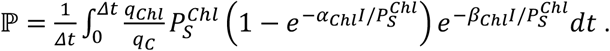

Here, following Platt et al.^45^: 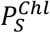 is an empirically constrained coefficient representing the Chlorophyll-a specific carbon fixation rate (*mol C* · (*mol Chl*)^−1^ · *s*^−1^) and 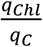 is the molar Chlorophyll-a to carbon ratio. *α_Chl_* and *β_Chl_* are empirically determined coefficients representing the initial slope of the photosynthesis-light relationship and photo-inhibition effects at high photon fluxes, respectively. Here we impose empirically determined values for *α_Chl_* and *β_Chl_* and 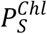 from the published study of Moore and Chisholm^24^. To find the maximum estimate for *Prochlorococcus* photosynthesis at different light intensities we use photo-physiological parameters for a High-Light adapted ecotype (MIT9215), acclimated at 70 *μmol* photons o *m*^−2^ · *s*^−1^ and a Low-Light adapted ecotype (MIT9211), acclimated 9 *μE.* Δt = 24 hours. *I* is the hourly PAR, estimated by scaling the observed noon value at each depth with a diurnal variation evaluated from astronomical formulae based on geographic location and time of year^33,34^. The Chlorophyll to Carbon ratio, 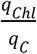, is estimated as a function of growth rate and light intensity using the model of Inomura et al.^46^ which was calibrated by laboratory data from Healey^47^.

The Chlorophyll to carbon ratio, 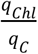, can be modeled as a function of growth rate and light intensity^46,48^. Here we use the Inomura^46^ model (equation 17 therein) where parameters were calibrated with laboratory data from Healey^47^. An initial guess of the growth rate and the empirically informed light intensity are used to estimate 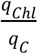, which is then used to evaluate the light-limited, photoautotrophic growth rate

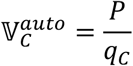

from which the Chlorophyll to carbon ratio is again updated. The light-limited growth rate is used to re-evaluate the Chlorophyll to carbon ratio. Repeating this sequence until the values converge, 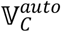 and 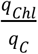 are solved iteratively.

The nitrogen-specific uptake rate of fixed nitrogen (*day*^−1^) is modeled as

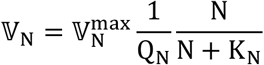

where values of the maximum uptake rate, 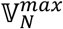 and half-saturation, *K_N_*, are determined from empirical allometric scalings^35^, along with a nitrogen cell quota *Q_N_* from Bertilsson et al.^37^ (0.77 *fmol N cell*^−1^).

The P-limited growth rate, or the phosphorus-specific uptake rate of phosphate (*day*^−1^), is modeled as

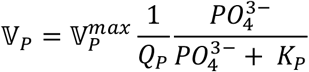

where values of the maximum uptake rate, 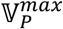 and half-saturation, *K_P_*, are determined from empirical allometric scalings^35^, along with a nitrogen cell quota *Q_P_* from Bertilsson et al.^37^ (0.048 *fmol P cell*^−1^).

Iron uptake is modeled as a linear function of cell surface area (*SA*), with rate constant 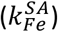 following Shaked et al.^36^.

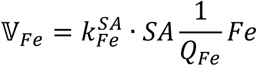

The potential light-, nitrogen-, phosphorus- and iron-limited growth rates 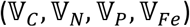 were evaluated at each depth in the water column and the minimum is the local modeled photo-autotrophic growth rate estimate, absent of mixotrophy (blue lines, Figure 2B, D). The the model is available at https://github.com/zhenwu0728/Prochlorococcus_Mixotrophy.

A significant premise of this study is that heterotrophy is providing for the shortfall in carbon under very low light conditions, but not nitrogen. It is known that Prochlorococcus can assimilate amino acids^9^ and therefore the stoichiometry of the heterotrophic contribution might alter the interpretations. However, it is also known that *Prochlorococcus* can exude amino acids^38^ which might cancel out the effects on the stoichiometry of *Prochlorococcus.*

For the estimates of photo-trophic growth rate from local environmental conditions (Figure 2) we employed photo-physiological parameters from laboratory cultures of *Prochlorococcus^24^.* For the purposes of this study, we have assumed that the photosynthetic rates predicted are Net Primary Production which means that autotrophic respiration has been accounted for in the measurement. However, the incubations in that study were of relatively short timescale (45 min), which might suggest they are perhaps more representative of Gross Primary Production. If this is the case, our estimates of photo-autotrophic would be even lower after accounting for autotrophic respiration, and thus would demand a higher contribution from heterotrophic carbon uptake. In this regard, our estimates might be considered a lower bound for organic carbon assimilation.

## Supporting information

Individual-based model simulation

## Acknowledgements

We thank the captain and crew of the R/V Mediterranean Explorer and Tom Reich, for help during the work at sea, Mike Krom and Anat Tsemel for the nutrient analyses, Maya Ofek-Lalzar for assistance with the bioinformatics analysis, Annett Grüttmüller for NanoSIMS routine operation, Ioannis Tsakalakis for help with hourly PAR estimation, and John Casey for the discussion about C uptakes rates. This study was supported by grant RGP0020/2016 from the Human Frontiers Science Program (to MV, HPG and DS) and by grant number 1635070/2016532 from the NSF-BSF program in Oceanography (NSFOCE-BSF, to DS). The NanoSIMS at the Leibnitz-Institute for Baltic Sea research in Warnemuende (IOW) was funded by the German Federal Ministry of Education and Research (BMBF), grant identifier 03F0626A. MJF and WZ are grateful for support from the Simons Foundation through the Simons Collaboration on Ocean Processes and Ecology (SCOPE 329108 to MJF) and the Simons Collaboration for Computational BIOgeochemical Modeling of marine EcosystemS (CBIOMES 549931 to MJF).

## Author contributions

DA, DRR, TLK, AV, MV and DS designed experiments, DRR, DA, TLK, LZ and DS performed experiments and field analyses, DRR, DA, TLK, AV, and FE performed NanoSIMS analyses, DA, DRR, TLK, AV, LZ, FE, HPG, MV and DS analyzed experimental results. WZ, MJF, OW and DS designed and executed the growth rate simulations. WZ designed and executed the individual-based model simulations. WZ, DA, DRR, TLK, MJF and DS wrote the manuscript with contributions from all authors.

